# Electrochemical Roughening and Carbon Nanotube Coating of Tetrodes for Chronic Single-Unit Recording

**DOI:** 10.1101/738245

**Authors:** Zifeng Xia, Gonzalo Arias-Gil, Martin Deckert, Maike Vollmer, Andrew Curran, Rodrigo Herrera-Molina, Marcel Brosch, Kristine Krug, Bertram Schmidt, Frank W. Ohl, Michael T. Lippert, Kentaroh Takagaki

## Abstract

Recording from single neurons in the brain for long periods of time has been a central goal in both basic neuroscience and translational neurology, in order to understand mechanisms underlying brain processes such as learning and to understand the pathogenesis of neurodynamic disease states ^1^. Recent advances in materials engineering, digital signal acquisition, and analysis algorithms have brought us closer to achieving this goal, and the possibility has gathered much public attention ^2,3^. However, it remains a challenge to record from the same units for weeks to months. Here, we record many high-quality tetrode neuronal signals reliably over long periods of time in both deep and superficial areas of the brain. We achieve this by combining electrochemical roughening and carbon nanotube coating of a flexible platinum/iridium substrate, with materials, packaging, and insertion optimized to minimize tip movement with brain pulsation. This “Magdeburger” probe enables recordings with long-term signal stability and high signal-to-noise ratio at a reasonable cost in both rodent brains and in substantially larger primate brains. Robust tetrode tracking of identified neurons over longer time periods, in multiple independently targeted areas of the brain, will allow fundamental advances in the study of cognitive learning, aging, and pathogenesis, and opens new possibilities for brain interfaces in humans.

---

Currently, four main classes of electrodes are standard for chronic *in vivo* recordings of neural activity: microwire arrays, Utah arrays, silicon probes, and flexible thin polyimide-based electrodes ^4^. These electrodes were designed to record from as many units (neurons) as possible—however, long-term stable recordings from tetrode-identified single units and juxtacellular recordings are rarely reported.

Microwire arrays are made of insulated sharpened metals, packaged in brush- or comb-like arrays ^5,6,7^. In selected cases, these electrodes demonstrate stable single unit recordings over months ^6,7^. Recently, solid-state-based versions of such a thin-wire brush approach have been proposed ^8,9,10^. The small diameter of these microwires minimizes tissue damage, but their flexibility in the transverse direction limits the accuracy of positioning individual electrode tips ^8,9,10^. Utah arrays, which are silicon-based microelectrode arrays with 96 contacts, are the most popular implant for chronic recordings in primates and humans and allow single-unit recordings and stimulation with high spatial resolution ^11,12^. However, they can only penetrate the cortex up to a depth of 1.5 mm ^13^ and are not suitable for recordings deep in the human brain or for individually-targeted multisite recordings. Furthermore, limited biocompatibility leads to encapsulation and cortical thinning after several months of implantation ^13,14,15,16,17^. With the previous approaches, each neuron is only recorded from a single microelectrode contact, and cannot benefit from the increased sorting precision afforded by multitrode sampling ^18,19,20^. Silicon shaft probes have addressed targeting of deeper structures in rodent brain and have packed many recording contacts, often into small areas ^21^. Hundreds of units can be recorded from a single insertion, but unit waveforms tend to “drift” over time and long-term recordings of stably identified units are seldom reported. Flexible thin-film probes can be implanted into nonsuperficial areas of the primate brain, and mitigate the problem of drift ^22,23,24^. Implantations can be scaled up using an automated sewing-machine like approach ^25^. However, the smoothness of these electrodes relative to their axial rigidity is not conducive to chronic anchoring. In summary, despite significant advances, current approaches do not reliably ensure chronic recordings from the same tetrode-identified units at various locations in the brain, especially in deeper areas of the primate brain.

The remaining challenges can be grouped conceptually into two classes: mechanical challenges and size challenges.

With respect to mechanical properties, a balance must be struck between two contradicting requirements: flexibility to allow pulsation together with brain tissue on the one hand, and rigidity to enable carrier-less targeted implantation on the other hand. Current approaches to striking a balance in this regard are to insert very soft electrodes via a thin carrier needle ^22,23^ or to inject them propulsively under pressure ^8,9,10,26^. These approaches reduce insertion trauma, but suffer from long-term instability and limited accuracy in targeting, respectively. Furthermore, injection-based approaches are currently optimized for superficial rodent cortex, and cannot reach deeper areas of the primate brain. We address this issue by choosing a direct implantation approach with axially rigid but transversely flexible soft Platinum-Iridium wire tetrodes. Recordings from classical twisted tetrode packages allow fundamentally more precise sorting of signal from different neighboring neurons, when compared to single contact recordings ^18,19^. Wires with several diameters were tested with/without heat curing. Implantation stability was screened with mock-surgical deep penetrations into semi-transparent 0.65% agarose gel, which is known to model the firmness of mammalian brain ^27^. Pt/Ir at 25 *μ*m was chosen due to its favorable mechanical handling during surgeries. Thinner Pt/Ir wires were too flexible for direct implantation, classical NiCr wires were too rigid for stability.

The second remaining challenge, with respect to electrode contact size, is to balance geometric compactness needed to isolate action potential signals from single neurons (which also implies higher impedance due to smaller contact area) and low impedance which is needed to record with high signal-to-noise ratio (due to reduced resistor noise). Low impedance recording with high signal-to-noise ratio is especially important in the awake cortex where local field potentials can manifest as high-amplitude fast spike-like signals, and low amplitude firing from distal units can easily be obscured by noise. Various materials and nanomaterials have been tested to lower impedance of microelectrode contacts. These include gold plating ^28,29^, carbon nanotube coating ^30,29^, and PEDOT coating ^31^. However, these coatings have not achieved wide adaption in neurophysiology, due to difficulties in the manufacturing process and problems with coating stability *in vivo*. Coating stability problems are particularly pronounced in packages such as sharp tungsten electrodes, Utah arrays, and side-contact silicon probe preparations, since axial advancement during implantation leads to shearing force in the delaminating direction. We addressed this challenge by developing a synergic electrochemical/carbon nanotube coating which leads to two-orders-of-magnitude impedance decrease relative to the native wire and one-order-of-magnitude decrease compared to state-of-the-art nanotube and other aforementioned coatings. This allowed us to minimize electrode noise (usually the limiting factor in such recordings) and to approach the amplifier noise levels of current neural amplifiers. As documented below, the synergic coating is resistant to aforementioned implantation shear.

For electrochemical roughening, tetrodes were immersed in 0.5M sulfuric acid solution and a train of electrical square wave pulses was applied for several minutes (see Methods and Supplementary Fig. 2). After roughening (Fig. 1a, lower left), sponginess at the sub-micron level is evident compared to the native cut surface (Fig. 1a, upper left). During the development of our protocol, we found that fine adjustment of roughening time and use of sonication to avoid gas buildup were key factors in order to prevent destructive changes of the electrode surface (Supplementary Figs. 1, 2). This roughening resulted in an electrode surface with lower impedances, well over an order of magnitude lower than the original cut electrodes (Fig. 1b), with slight improvement of charge transfer capacity (Fig. 1c). Subsequent carbon nanotube electroplating of the roughened surface lead to a further decrease in electrode impedance to approximately two orders of magnitude of native wire (Fig. 1b), and further improved charge transfer capacity (see Fig. 1c and Methods). Impedance spectroscopy showed log-linear decline over the range of neuronal-spike-relevant frequencies, indicating a predominantly capacitative electrode interface (Supplementary Fig. 4). Acute implantation of these ultra-low impedance flexible electrodes lead to recording of excellent single-trial unit signal in both rodent and primate brain (Fig. 2).

**Figure 1.**
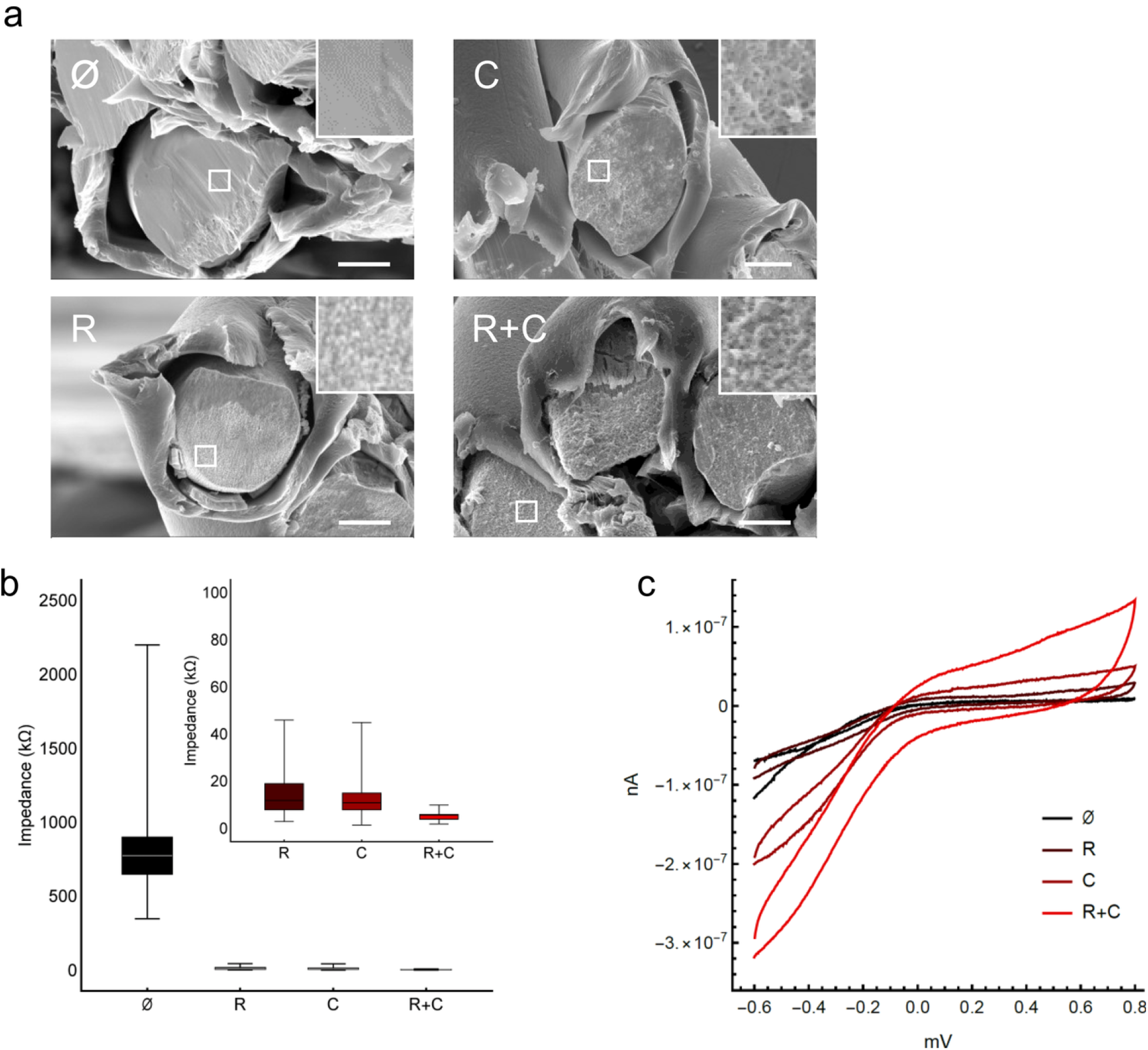
Characterization and comparison of various electrode surfaces. **a**, Electron-microscope images of tetrode contacts with variable surface processing: cut/untreated (∅, top-left), coated with carbon nanotubes (CNTs) (C, top-right), electrochemically roughened (R, bottom-left), and electrochemically roughened and CNT coated (R+C, bottom-right). Scale bars: 10 μm, insets 5x magnification. **b**, Impedances of tetrode contacts with various surface processing. Both CNT coated (8-15 kOhm, upper and lower quartiles) and electrochemically roughened (8-19 kOhm) contacts have impedance reduced by almost two orders of magnitude compared to untreated contacts cut with sharp scissors (650-900 kOhm). Contacts which are electrochemically roughened and coated with CNTs achieve even lower impedance (4-6 kOhm). **c**, Cyclic voltammetry scan of differently processed tetrodes. Both CNT coating and roughening increase the charge transfer across the contact surface. Even higher charge transfer increase can be achieved by combining coating and roughening.

**Figure 2.**
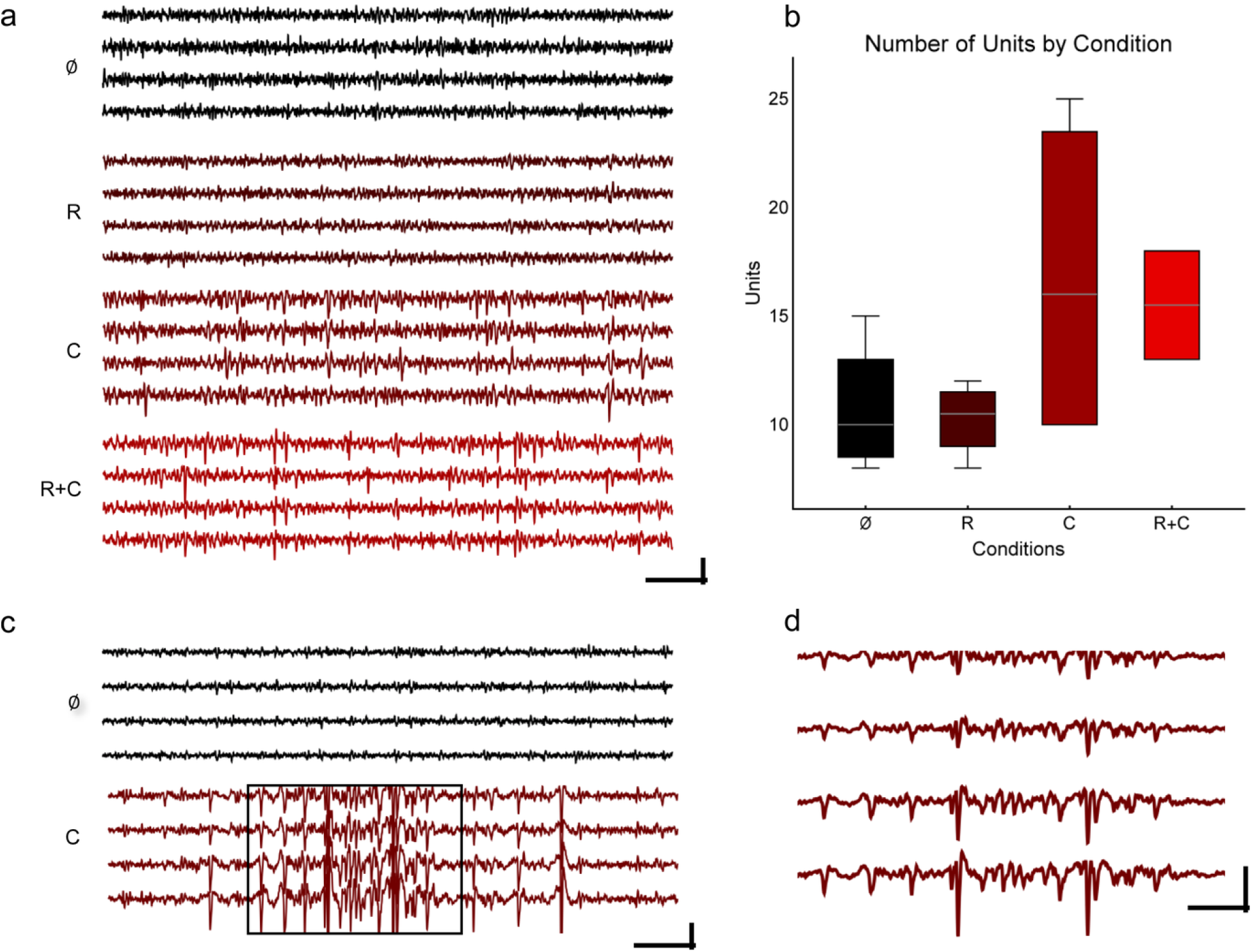
Effect of roughening and carbon nanotube coating on neuronal signals. **a**, Example traces from acute recordings in the globus pallidus externus of rat, with four different conditions (scale bar: 10 ms/100 μV). Although both cut and roughened contacts do show some unit spiking, the signal-to-noise ratio is not favorable. Clearer unit signals with better signal-to-noise ratio are seen with coated tetrode preparations. Abbreviations for the conditions are as described in the legend to Fig. 1a. **b**, Number of units detected in acute globus pallidus externus recordings per tetrode, by condition. Recordings were made acutely after implantation. A clear benefit in unit count is seen associated with carbon nanotube coating. Boxes denote quartiles, whiskers denote most extreme outliers. **c**, Representative acute traces recorded from area 46 in a rhesus macaque, anesthetized with isoflurane by inhalation and augmented with sufentanil citrate (i.v.) (scale bar: 10 ms/100 μV). Data are chopped to display low-amplitude activity, see d for full scale. **d**, Boxed segment in c, at lower magnification (scale bar: 10 ms/250 μV). Clear juxtacellular recordings of a bursting unit, approximately 500 μV in amplitude, are seen. This excellent signal demonstrates the potential for translation to primate cortex, including for translational human use.

Of note, initial ramping of coating voltage proved to be critical (Supplementary Fig. 3): ramping did not change electron-micrographic surface appearance or ohmic characteristics, but prevented crosslinking and lead to overall improvement in coating stability during and immediately after implantation (Supplementary Fig. 3, Supplementary Table 5).

Packaging and surgical approach are also important to achieve atraumatic implantation and stable chronic single unit recordings. The most successful approaches to date for recording from clearly isolated single units while maintaining stable waveforms over several weeks have been based on thin and flexible microwires ^6,32,8,7,26^. This success is presumably because the thin microwires move flexibly with brain micromovements, thereby keeping the tip in a stable position relative to the units it is recording from. Moreover, microwires can be inserted without a carrier, thus, reducing the risk of tissue damage. Penetration into deep areas remains a fundamental issue, however, especially for silicon-based propulsive injection approaches.^8,26^

Recently, tetrodes have regained interest for better chronic unit isolation, since the four independent channels allow fundamentally more precise sorting and provide a much better basis to judge unit stability in the context of drifting waveforms ^20,33^. Our data support this approach. In order to achieve our carrier-less surgical approach, we designed a 3D-printed polylactic acid tetrode holder which holds the tetrodes with the necessary stability to allow axial insertion without transverse deformation or bending (Fig. 3e). Anatomically, the rodent skull is significantly more plastic than the primate skull, often with open cranial sutures well into adulthood. This poses a difficulty when mounting electrode assemblies for chronic recording, because conventional high-channel-count connectors (such as those available from Omnetics) are designed to have particularly high stability, and therefore require non-negligible insertion force. A typical neurophysiological plug package, mounted perpendicularly, applies perpendicular force to the dorsal surface of a rodent skull leading to dorsoventral skull/brain deformation during each connection cycle. To avoid this, we have designed a laterally-oriented low-insertion-force Hirose board-to-board plug and tetrode holder package (Fig. 3b, c). The plug is stabilized by a custom 3-D printed clip. Another advantage of this connector strategy is that it is cost-effective, with the Hirose plugs available in the 10-20 cent range (compared to the 100 euro range for conventional plugs). This package should also allow effective mounting of other electrode types on the rodent skull.

**Figure 3.**
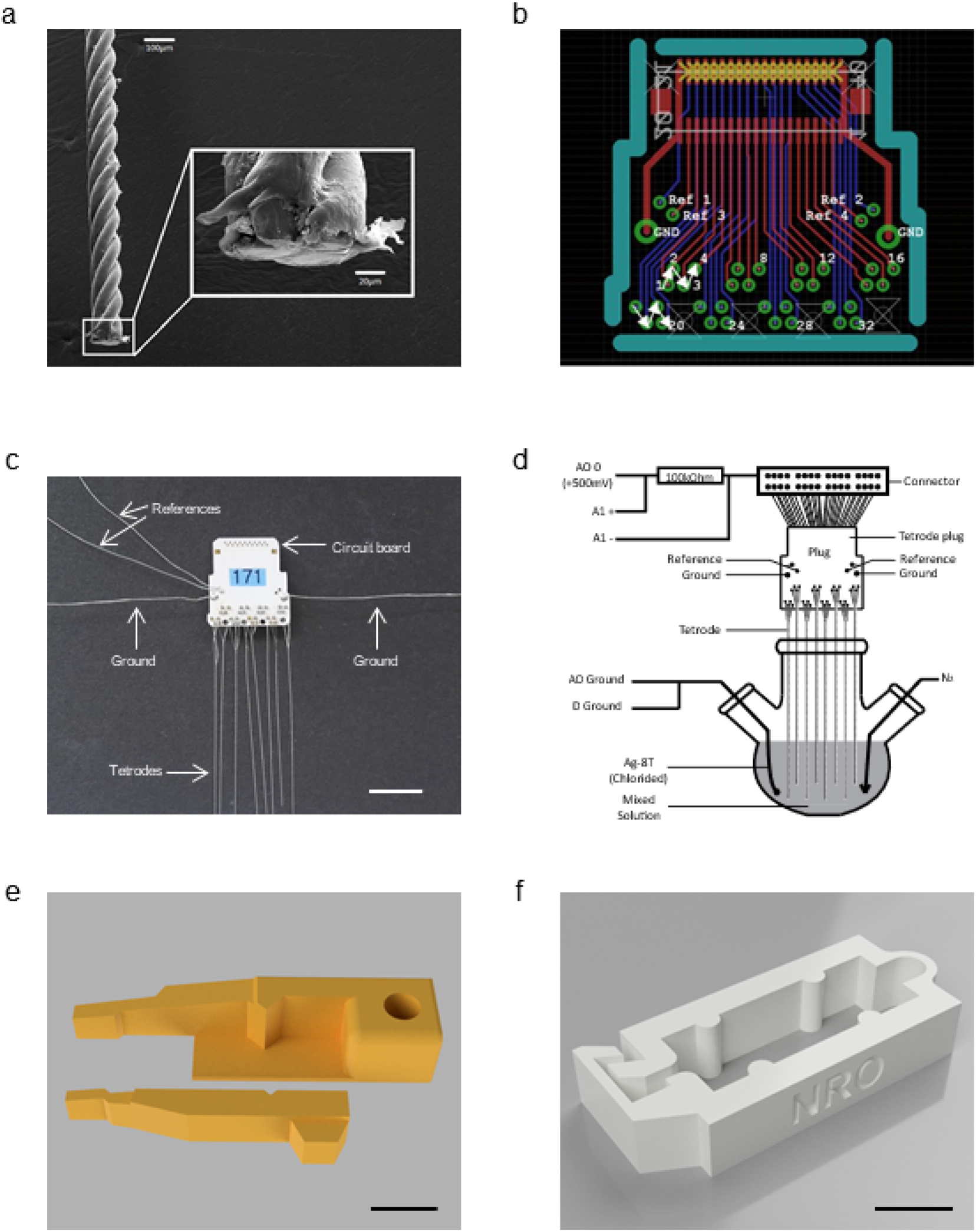
Technical setup and engineering required for manufacture and low-trauma implantation of roughened/coated tetrodes. **a**, Electron micrograph of a micromanufactured twisted tetrode, polished with a circular grinder to 90 degrees at the tip. **b**, A custom circuit board serves as an implantable contact interface for flexible tetrodes. It has 32 gold plated contact holes for eight tetrodes, 4 gold plated holes for reference electrodes, and 2 gold plated holes for ground electrodes. The circuit board is mounted perpendicularly to the dorsal surface of the skull, allowing insertion to be in the lateral direction (which does not compress the skull downwards). This design uses a low-cost low-insertion-force Hirose DF12 board-to-board connector. **c**, Sample implant with tetrode circuit board (b.), ground electrodes, chlorided reference electrodes, and 8 roughened and carbon nanotube coated tetrodes mounted with gold pins (scale bar: 10 mm). **d**, Experimental setup for tetrode carbon nanotube coating experimental setup. A voltage of +550 mV is applied to each electrode individually. **e**, A 3D-printed custom electrode holder is used for holding/driving tetrodes during implantation. Scale bar: 5 mm. **f**, A 3D-printed clip, which allows precise fixation of the plug connecting implant and headstage. This is important due to the lower insertion force of the board-to-board plugs used (scale bar: 5 mm).

Carbon nanotube *in vivo* electrodes have not been commercialized and have been used rarely to date ^30,29,34^, primarily because of three factors: fickleness of the coating process, instability of prior coatings, easy shearing (discussed above), and shelf life and stability in solution (Sherman Wiebe, personal communication). We evaluated the shelf life of our coating over the course of one year, by conducting longitudinal impedance measurements in 3M KCl solution. We found that our coating maintains stable impedances for over one year, which is compatible with commercialization of batch-manufactured lots (Fig. 4a). It is also of interest that the impedance of the electrode contacts themselves are significantly more stable than the impedance of unwanted crosslinks within tetrode contacts, which increased over one order of magnitude over the same year of measurement (Fig. 4b). As unwanted crosslinks tend to rise in impedance during shelf storage, coating protocols for mass production can be biased toward overcoating and lowest possible impedances (which inevitably leads to accidental crosslinking).

**Figure 4.**
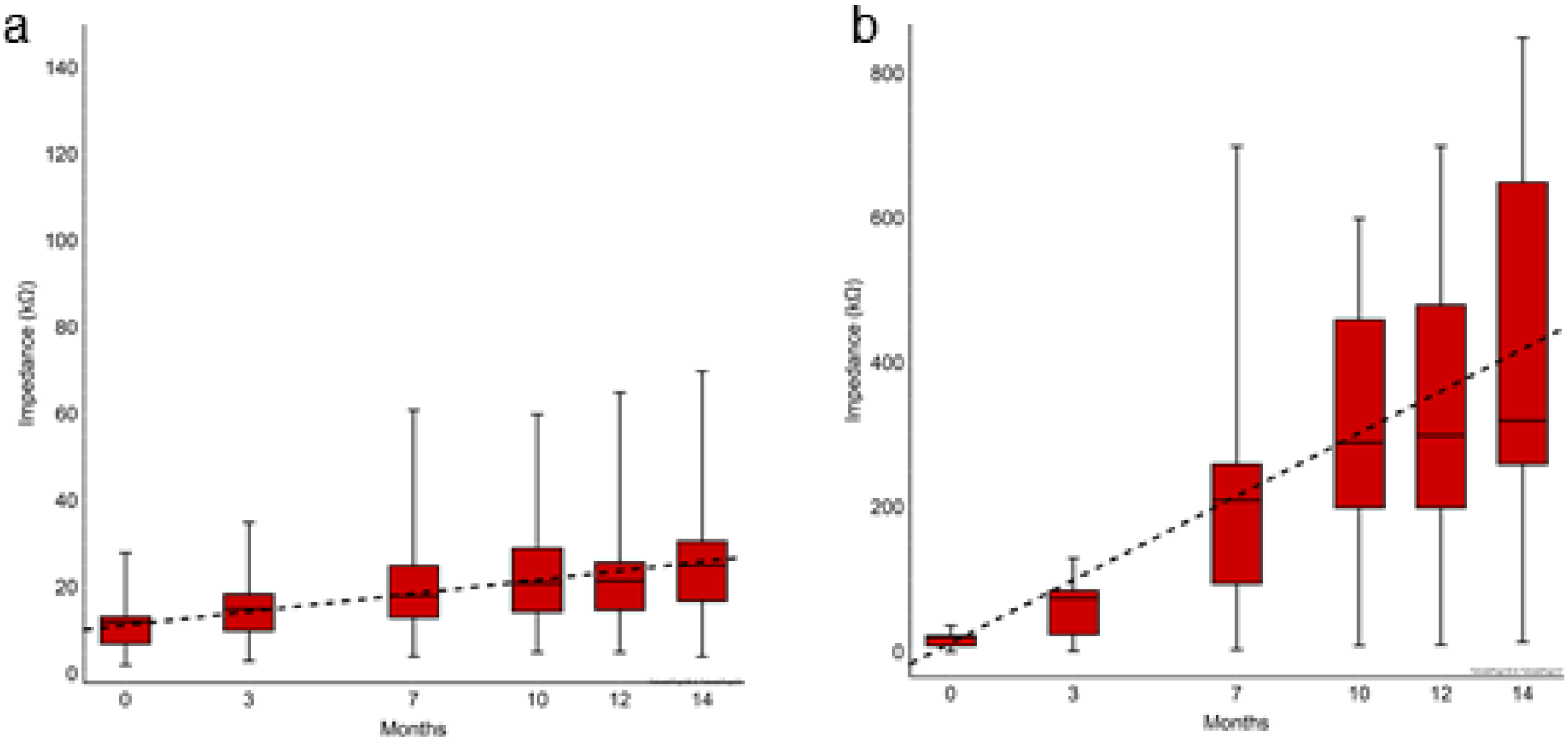
Shelf life of carbon nanotube coating. **a**, Shelf life of carbon nanotube coating in air. An approximate doubling of impedance is seen in 1 year, which is fully compatible with commercialization. Eight tetrode contacts per condition were measured serially for each graph. Boxes denote quartiles, whiskers denote most extreme outliers. **b**, Impedances of unwanted carbon nanotube crosslinks (between individual contacts within a tetrode) over time. These crosslinks are prevented by ramping of coating currents (Supplementary Fig. 3). As seen, unwanted crosslinks increase their impedances much more rapidly than the main contact (a), probably due to oxidation, since the crosslink bridges are thinner and more exposed to air. Since unwanted low-impedance crosslinks are self-limited in this manner, one can tolerate a bias toward excessive coating and crosslink formation for production scale-up and commercialization.

In the past, several groups have reported single unit recordings from identified neurons lasting for months, especially in primates ^6,35,7,20^. Other groups have reported long-lasting unit recordings from rodents as well, with some success ^32^. However, no group to date has been able to record from a large number of clearly identifiable tetrode units and juxtacellular recordings for extended periods of time. Using our “Magdeburger” tetrodes, we were able to stably record from a large variety of areas in the rodent and monkey brain with large signal-to-noise ratios (Fig. 2a, c-d, Fig. 5g). For validation purposes, we focused here particularly on signals from juxtacellular recordings. Juxtacellular recordings are defined as unit recordings with very large amplitude waveforms recorded from neurons, which are presumably adjacent (“juxta”) to the electrode tip ^36^. The regular occurence and stability of these juxtacellular recordings in our preparation (Fig. 5) is remarkable, because we did not intentionally target them (the usual procedure for juxtacellular recordings with glass electrodes) ^36^. Instead, we believe that our juxtacellular signal develops over the course of several weeks of post-surgical recovery; the biocompatibility of our coating was permissive to outgrowth and direct contact of the electrode contacts by neurons (Supplementary Fig. 9), resulting in fortuitous juxtacellular arrangements arising several weeks after implantation (Fig. 5). This juxtacellular effect allows signal-to-noise ratios approaching those of intracellular recordings, with a less invasive extracellular configuration (Fig. 5)^36^. In our case, this juxtacellular effect can be stable for weeks instead of the usual hours (Fig. 5). We have yet to see any cases of post-mortem neuropathology in our successful long-term recordings such as implantation tracts, gliosis, or atrophy; this is also supportive of the contention that our biomicroengineering and material engineering approaches are enabling long-term biocompatibility.

**Figure 5.**
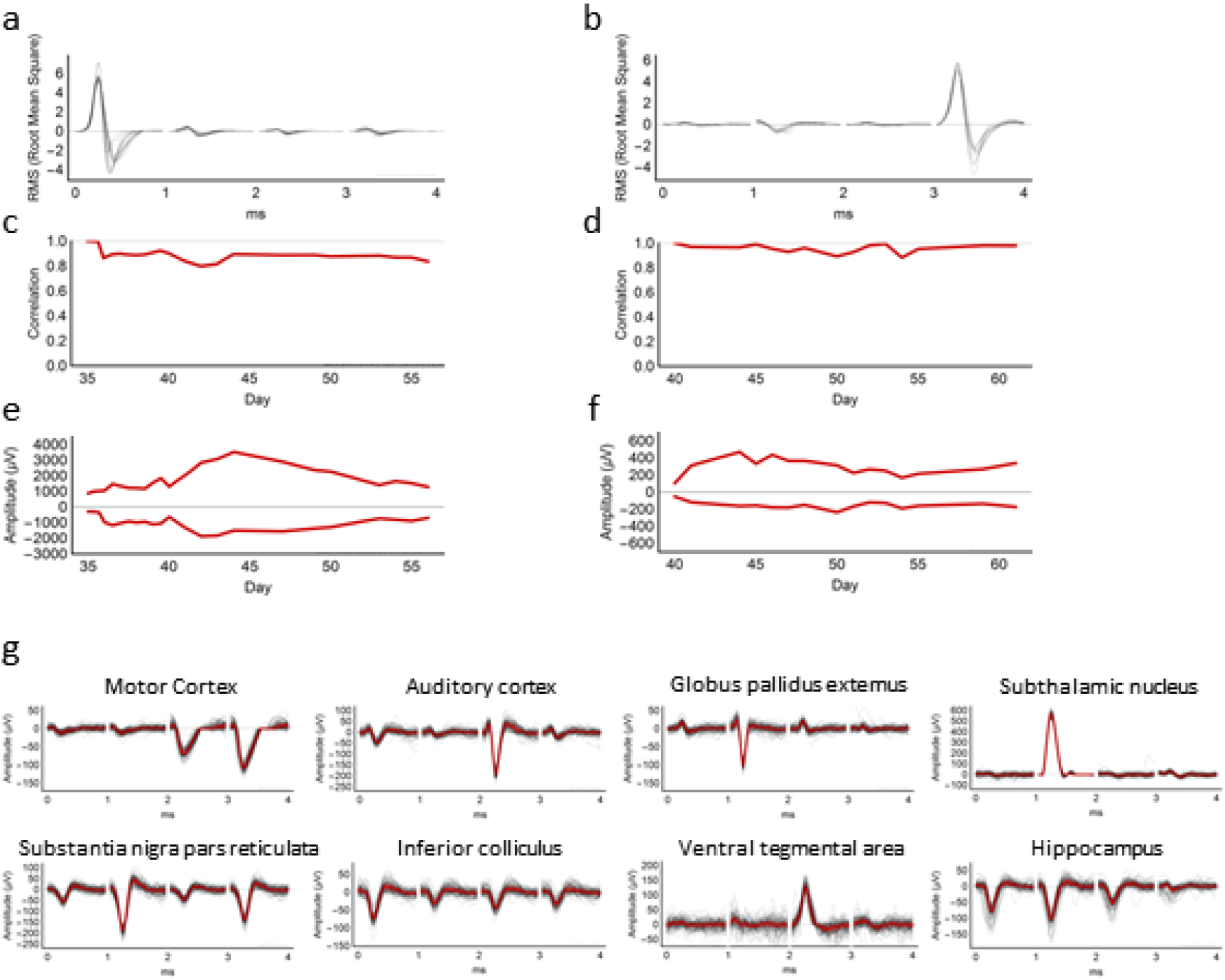
Large single units and “juxtacellular” waveforms can be recorded over weeks. **a**, **b**, Stable juxtacellular unit waveforms recorded daily over weeks (with breaks for experimenter recovery), in rat. Each black line is a waveform normalized each day to its mean amplitude (1), with waveforms from all four tetrodes shown sequentially. This overlay shows that overall waveform shapes are stable over 3 weeks, in each case. These waveforms are displayed without normalization in Supplementary Fig. 8. **c**, **d**, Waveform shapes are stable over time (days after implantation), as revealed by Pearson correlation coefficients of the data in a and b, relative to the waveform on the first plotted day. The maintenance of overall waveform over time suggests a biological impedance change, rather than a geometric location change of the electrode, as the cause of our minimal drift. **e**, **f**, Waveform peaks and troughs of the same units wax and wane over the course of weeks. The maximum and minimum of the data in a and b were plotted as a timecourse. Note that the maximum waveform amplitude reaches well into the mV range, which is remarkable for unit recordings. The slow time course of changes further suggests a biological process as the cause of waveform drift, rather than mechanical movement of the brain or electrode tip, which would lead to sudden changes in daily recordings. **g**, Such large units can be recorded from all over the rodent brain, including areas not reachable by current silicon-based approaches. Black waveforms are 100 superimposed filtered waveforms, extracted randomly from all waveforms matching each unit. Red lines depict the median of these black waveforms.

In summary, we have applied two material modifications to the classical twisted tetrode. First, we selected a soft metal substrate to make our tetrodes as flexible as possible in the transverse direction and implanted directly without a carrier or a head-mounted electrode drive. Despite the flexibility of our electrodes, the twisted braids provide contour to the electrode shank, allowing stable anchoring within the tissue. Second, we developed an electrochemical roughening procedure which improves the microscopic surface characteristics of our cut electrode surface, and a synergic carbon nanotube coating procedure to coat the roughened surface. The result is a surface which is fractal at the molecular-scale but nonetheless mechanically stable during implantation, and featuring an impedance reduction by two orders of magnitude to the sub 10 kOhm range with large charge transfer capacity. Our coating is also biologically stable, (Supplementary Fig. 9), allowing proximal neurite outgrowth and juxtacellular unit recordings of unprecedented quality. We have yet to observe any long-term neuropathological changes, which is quite atypical for such implants but is a necessary condition for long-term experimental and clinical use. With this "Magdeburger"tetrode, we are able to record stably at juxtacellular quality over weeks instead of just hours, which is also unprecedented. Our package allows independent targeting of multiple areas, which is not possible in current chronic recording approaches. These advances will allow qualitatively new approaches in basic neurophysiology research, in translational neurology, and potentially in future human brain-computer interfaces.

## Online Methods

### Tetrode Manufacture and Roughening

A base material of platinum 20% iridium wire was used for tetrode manufacture (California Fine Wire Company, USA; teflon coated and stress relieved with 31.75 μm/25 μm shielded/metal diameter). These wires were twisted (without heat curing) and mounted on a custom-designed circuit board, to create an implant with 8 tetrodes (32 recording channels; Supplemental Figure). The implant also included 2 ground wires (bare silver wire Ag-8W, 99.99%, 200μm diameter, Science Products GmbH) to be attached to skull screws and 2 reference electrodes (teflon-coated silver wire Ag-5T, 99.99%, 125 μm/200 μm metal/coated diameters, balled and chlorided), which were implanted superficially on neurophysiologically neutral sites such as cerebellum and parietal cortex.

Tetrodes were cut at a 45 degree angle just prior to roughening, using sharp carbide scissors (Fine Science Tools GmbH, Heidelberg Germany). The sharpness of these scissors and the angle of the cut may have a slight effect on final impedances (although not clearly significant). Although not systematically tested on large numbers of probes, polishing of tetrode tips to provide a smooth surface (Supplemental Fig. 7) seems to increase tetrode impedances, perhaps due to the microscopically rough cut surface contributing to the stability of our carbon nanotube coat.

Each tetrode was electrochemically roughened with a protocol modified from previous reports on different metals ^36^. First, the cut surface was immersed in 0.5M sulfuric acid solution and roughened by applying a train of square waves at 1 kHz with 50% duty cycle for 60 seconds against a Ag|Ag2SO4 reference electrode. The upper and lower potentials of the square wave were +2.4 V and −0.4 V respectively. The effective duration of roughening time was much shorter than reported previously for platinum ^37^. Of note, a sonicator was used to prevent gas formation at the tip of tetrodes during roughening. Without this, strange oxidization artifacts were seen, which featured higher impedances (Supplemental Fig. 1). Tetrodes were maintained in the sulfuric acid solution at −0.4 V for 3 minutes to reduce oxides prior to rinsing with deionized water and drying.

### Carbon nanotube coating

Each tetrode was coated with carbon nanotubes, using a protocol modified from prior reports ^38^. The reagents used for CNT coating were as follows: carbon nanotube powder (NC3151, research grade – short thin MWCNT 95+%C purity and surface modified COOH, Nanocyl); poly (sodium 4-styrenesulfate) (average Mw ~70,000, powder, product #243051, Sigma-Aldrich); pyrrole (reagent grade 98%, product #131709, Sigma-Aldrich).

First, 50 mg of carbon nanotube powder was added to 50 ml of distilled water and placed in a beaker in an ice bath. The carbon nanotube solution was then thoroughly sonicated with a horn sonicator for 45-60 min and transferred to a cylindrical flask. The solution was stirred gently on ice with a magnetic stirrer, while *N*_2_ was gently bubbled to eliminate oxygen from the solution and minimize oxidation. Next, 200mg of poly (Na 4-styrenesulfate) was added and mixed well with a magnetic stirrer until dissolved. Finally, 1.736 ml of pyrrole was added. This solution provides stable coating results for at least 120 min after Pyrrole is added to the preparation.

Eight tetrode tips cut at 45° were immersed in the solution for coating. Prior to coating, these tetrodes were either freshly cut with sharp carbide scissors, or roughened with the procedure outlined above. A voltage of +550 mV for a total coating time of 2100 ms (including linear ramp times of 50ms) was applied to each electrode. Applying voltage to multiple tetrode contacts simultaneously was avoided, since it lead to crosslinking (Supplemental Fig.3).

### Impedance Measurements, Impedance Spectroscopy, and Cyclic Voltammetry

Impedance measurements, impedance spectroscopy, and cyclic voltammetry were conducted independently on several instruments in 3M KCl solution against Ag/AgCl. Standard serial measurements were made at 1 kHz with the Metal Electrode Impedance Tester IMP-1 (Bak Electronics, Umatilla, USA). Impedance spectroscopy was done with nanoZ model 1.2 and nanoZ software version 1.4.0 (White Matter LLC, Seattle USA). Fifteen measurement cycles were acquired at test frequencies from 2Hz to 2kHz. Additional impedance spectroscopy and cyclic voltammetry were performed on a Interface 1010E Potentiostat (Gamry Intruments, Warminster, USA) against a dual-diaphragm Ag/AgCl reference electrode filed with 3M KCl.

### Electrode Implantation and Neural Recordings

Rodent electrode implantation was performed under standard anesthetic conditions with 50 mg/kg pentobarbital injected intraperitoneally. Under stereotaxic and physiologic guidance, electrodes were slowly advanced through a burr hole in the skull using the holder shown in Figure 3e until robust units characteristic of each given target area were seen. Electrodes were cemented at their insertion point to the skull, using UV-curing cyanoacrylate glue (TDS LOCTITE® 4305, Henkel AG & Co. KGaA, Dusseldorf, Germany) and Resilit-S Cold-curing acrylic resin (Erkodent(r) Erich Kopp GmbH, Pfalzgrafenweiler, Germany). Surgical procedures were approved by the state of Saxony-Anhalt, Saxony.

Monkey recordings were obtained from one adult male macaque monkey under combined isoflurane (0.5-1.0%)/sufentanil citrate (from 0.6 lg/kg/h i.v.) anesthesia in the context of a terminal acute experiment. For details of the set up and procedures, see Ahmed, 2012 ^39^. Housing and husbandry were in compliance with the ARRIVE guidelines of the European Directive (2010/63/EU) for the care and use of laboratory animals. All animal procedures on monkeys were carried out in accordance with Home Office (UK) Regulations and European Union guidelines (EU directive 86/609/EEC; EU Directive 2010/63/EU).

Neural recordings were conducted with Digital Lynx 4SX (Neuralynx, Dublin, Ireland) in a Faraday cage inside an acoustic chamber. Recordings were pre-filtered at 10 to 9000 Hz, and digitized at 14-bits. Rodent recordings were commutated using a custom mechanical swivel (Takagaki et al., in preparation), and rodents were recorded from under freely moving awake-behaving conditions.

Analysis was conducted using Scala/Java (EPFL/Oracle) and Wolfram Mathematica 11.3.0.0 and 12.0 (Wolfram Research, United Kingdom), using block-based modular data stream processing as reported previously ^39^.

### Cell culture, immunocytochemistry, and cell imaging

Primary cultures of rat hippocampal neurons were obtained as previously described ^41,42^. After one week in culture on 12 mm glass coverslips, neurons were exposed to 5 μg/ml carbon nanotubes for 10 days.

Hippocampal neurons were fixed with 4% PFA for 8 mins and then gently incubated with PBS 1X for 2 mins and then with a solution containing 10% horse serum, 0.1 mM glycine, and 0.1% Triton X-100 in Hanks’ balanced salt solution two times for 5 min. Pyramidal neurons were morphologically identified based on the side and shape of cell body as observed using anti-MAP2 guinea pig (Synaptic Systems GmbH, Göttingen, Germany; 1:1,000). To visualize synaptic contacts, samples were incubated with rabbit polyclonal synaptophysin 1 (Synaptic Systems; 1:500) for 2 hours at 4°C. Subsequently, samples were incubated with anti-rabbit Cy3-conjugated donkey secondary antibody (1:1,000) for 1 h. Then samples were gently washed and mounted with Mowiol. Fluorescence was visualized using a Zeiss AXIO Imager A2 microscope equipped with a CCD camera (Visitron Systems; camera binning = 1, pixel size = 0.10238 × 0.10238 μm, pixel depth = 16 bytes) using a X63 (1.4 NA) objective.

## Acknowledgements

The authors would like to thank Silvia Vieweg and Beate Traore for excellent technical and administrative support. This work was supported by the Leibniz Institute for Neurobiology, Priority Program 1665 of the DFG, the Alexander von Humboldt Foundation, the JMilk Foundation, and the Wellcome Trust (101092/Z/13/Z).

## Author contributions

This study was conceived by KT and MTL. Experiments were planned and executed by ZX, KT, GAG, MD, MB, MV, AC, RHM, MTL, and KK. Figures were created by ZX, KT, MD, and MB. Expertise and manuscript writing/editing was provided by KT, MTL, ZX, GAG, MV, AC, RHM, BS, and FWO, and KK. Correspondence and material requests should be addressed to KT.

## Supplementary Information

**Supplementary Figure 1.**
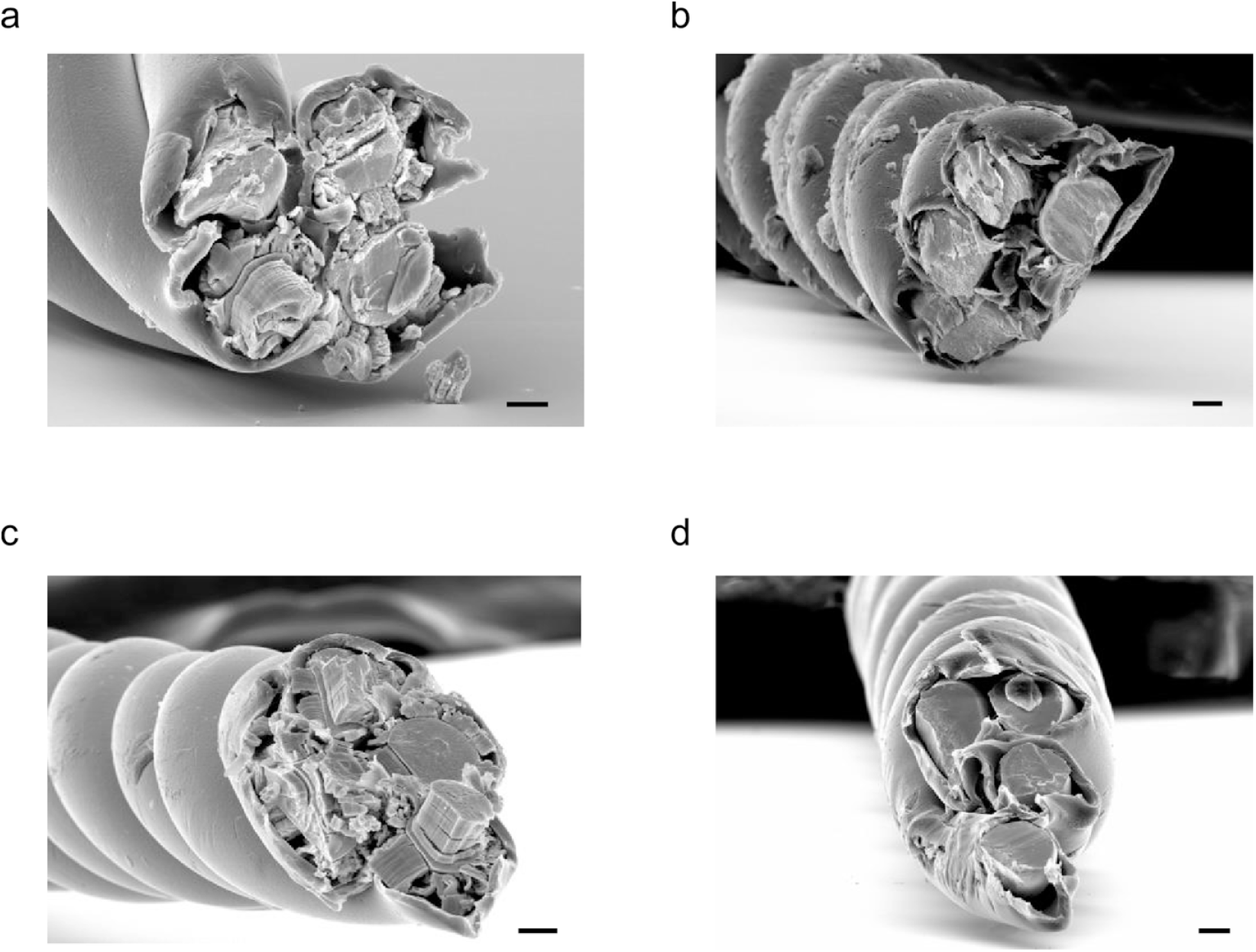
Abnormal oxidation products. Abnormal “tree-barking” oxidation products were observed during electrochemical roughening, prior to optimization of our protocols. This is presumably due to gas emission and cracking. Scale bars: 10 μm. **a**, Each contact electrochemically pulsed with square wave potentials at +2.4 V and −0.4 V for 3 minutes, without sonication. **b**, All four contacts electrochemically pulsed simultaneously for 55 seconds, and kept in roughening H2SO4 solution at −0.4 V for 30 seconds at the end, without sonication. **c**, Electrochemical pulsing with square wave potentials at +2.4 V and +0.4 V for 3 minutes, without sonication. **d**, Electrochemical pulsing with square wave potentials at −0.4 V and −2.4 V for 3 minutes, without sonication.

**Supplementary Figure 2.**
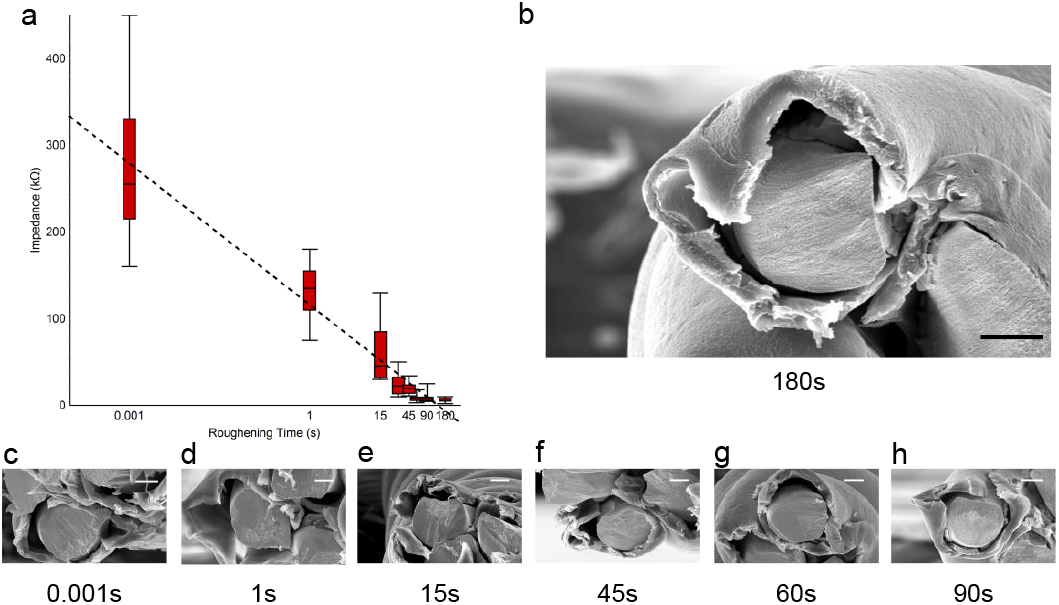
Tetrode impedance drops with roughening time. **a**, The cut surface of a tetrode was immersed and electrochemically roughened in sulfuric acid solution with variable pulsing durations. The log-linear fit indicated by the dotted line (115.934 −54.3555 log10t) indicates an exponential process: longer electrochemical roughening results in lower impedance. At roughening times >= 60s, the impedance decrease saturates at about 8kOhm (t in seconds). **b**, Electron micrographs of a tetrode contact, with saturated electrochemical roughening (180 s pulsation). Scale bar: 10 μm. **c-h**, Electron micrographs of representative tetrode contacts, roughening with different pulsation times as plotted in **a**. Scale bars: 10 μm.

**Supplementary Figure 3.**
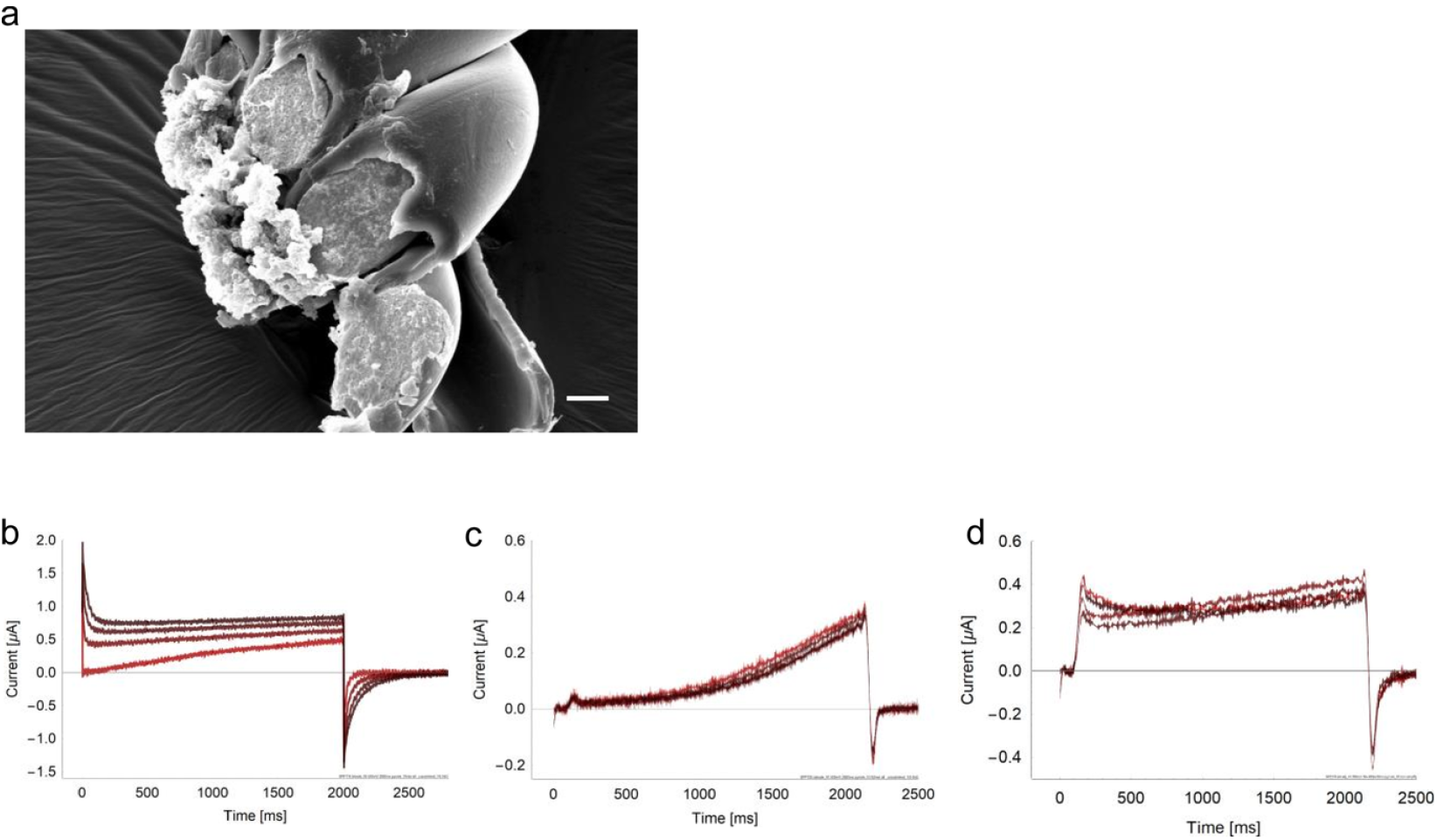
Crosslinks due to CNT overcoating. **a**, Crosslinks within a tetrode can be eliminated by applying a high voltage (e.g. approx. 18 V) across the two crosslinked contacts. Empirically, this seemed to increase the incidence of carbon nanotubes falling off during or after implantation (Supplementary Table 5). Scale bar: 10 μm. Boxes denote quartiles, whiskers denote most extreme outliers. **b-d**, Ramping of coating current prevents crosslinking. **b**, Sample coating currents without ramp. When coating each tetrode contact one by one, the 2nd, 3rd, and 4th contacts (darker colors) often show a higher starting point already at the very beginning of voltage application, which demonstrates that coating of previous contacts affect subsequent contacts. This may be due to capacitive currents leading to poorly controlled CNT coating and crosslinks at non-targeted contacts. **c**, Sample coating currents with 50 ms ramp added to applied voltage. Adding a ramp prevented crosslinks almost completely, and initial capacitive currents are not seen. **d**, Sample coating currents for CNT coating of a roughened tetrode. All traces have high starting points, due to the higher capacitance of roughened contacts, but this did not lead to spurious crosslinking.

**Supplementary Figure 4.**
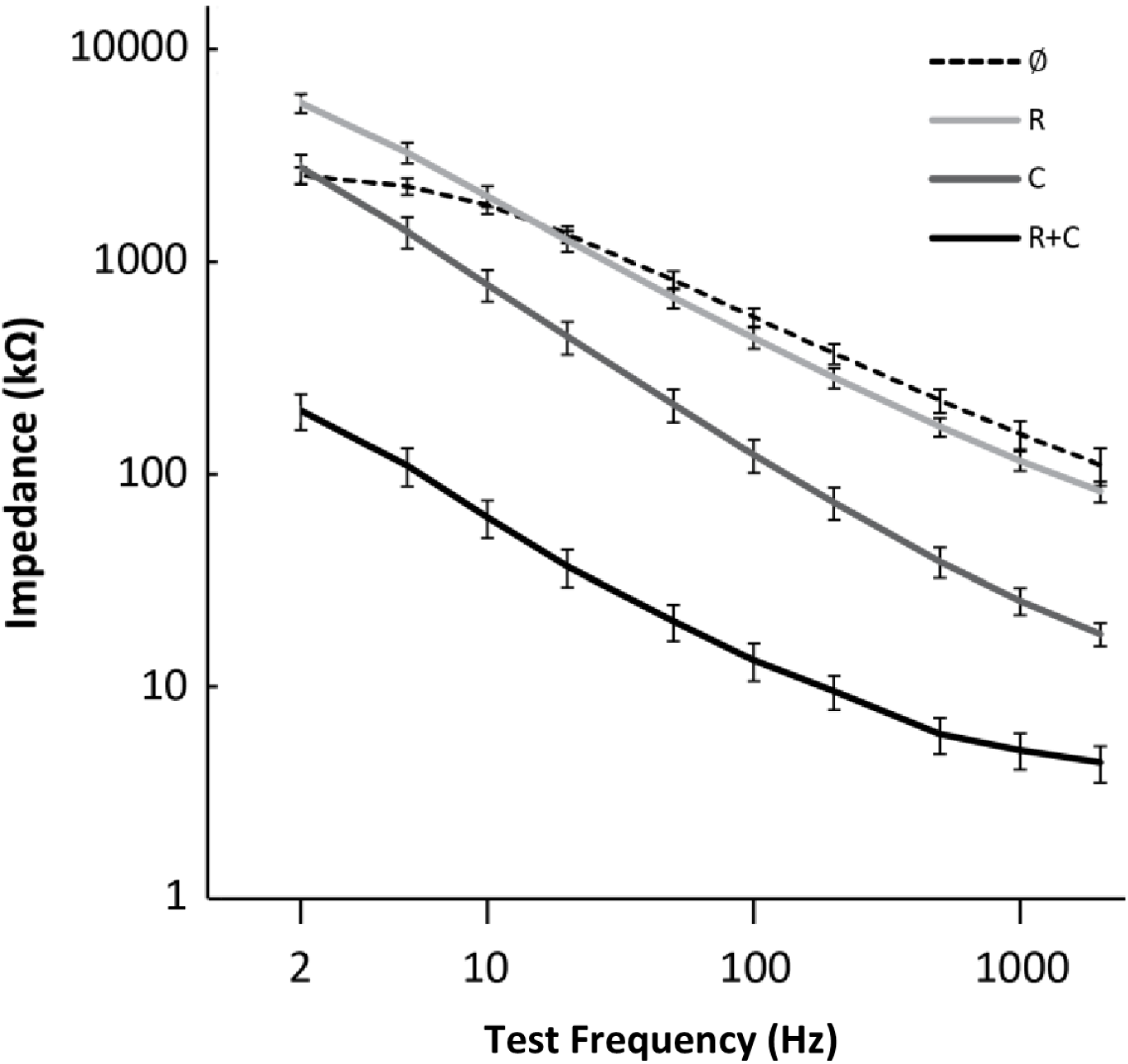
Impedance Spectroscopy. Electrodes with all types of finish show log-linear reduction in impedance over frequency, as expected from a mostly capacitive electrode interface. While cut wires show the highest impedances, coated as well as roughened and coated electrodes have low impedances optimal for low-noise recording of neuronal signals.

**Supplementary Table 5.** Coating failure during the first week of implantation. Loss of carbon nanotube coating in single channels of a tetrode occurs exclusively during the first week after implantation, even when the coating survives the acute implantation surgery with no change in signal characteristics or impedance. Given the time course, it is likely that an active biological process such as glial phagocytosis is involved.

| **Supplementary Table 5 \| Chances of CNT loss during or within one week after implantation** |  |
| --- | --- |
|  | Total number of contacts |
| CNT coating | 3.30% (39/1182) |
| Roughening & CNT coating | 0.34% (2/587) |

**Supplementary Figure 6.**
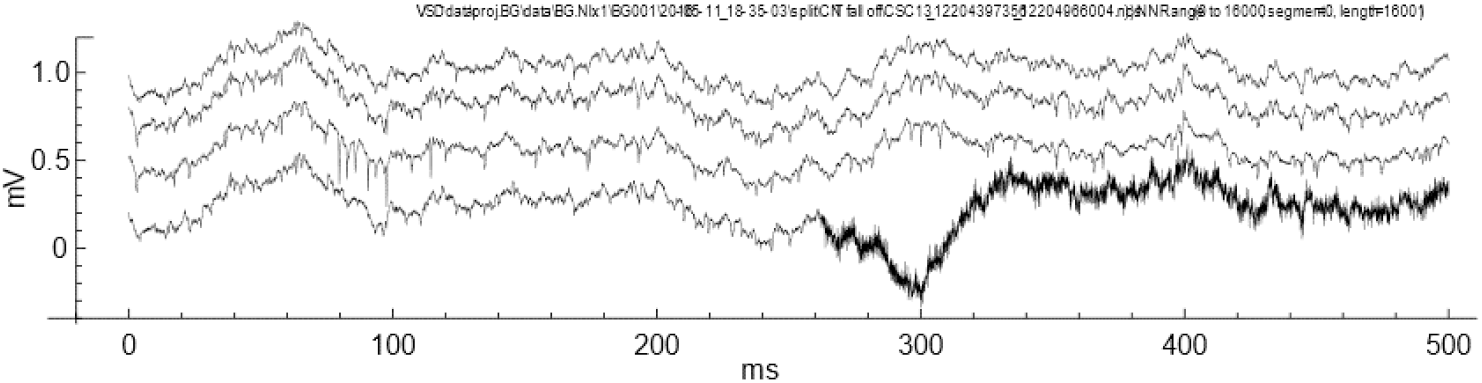
Carbon nanotube coating falling off during implantation. Recording during an implantation operation with a carbon nanotube coated tetrode. Sudden increase of baseline noise was seen on one channel, which we interpret as either loss of carbon nanotube coating or disrupted contact between carbon nanotubes and metal surface. This serves to demonstrate the signal effects of our coating. Of note is also that our high capacitance electrode interface also allows simultaneous recording of slow and fast local field potentials.

**Supplementary Figure 7.**
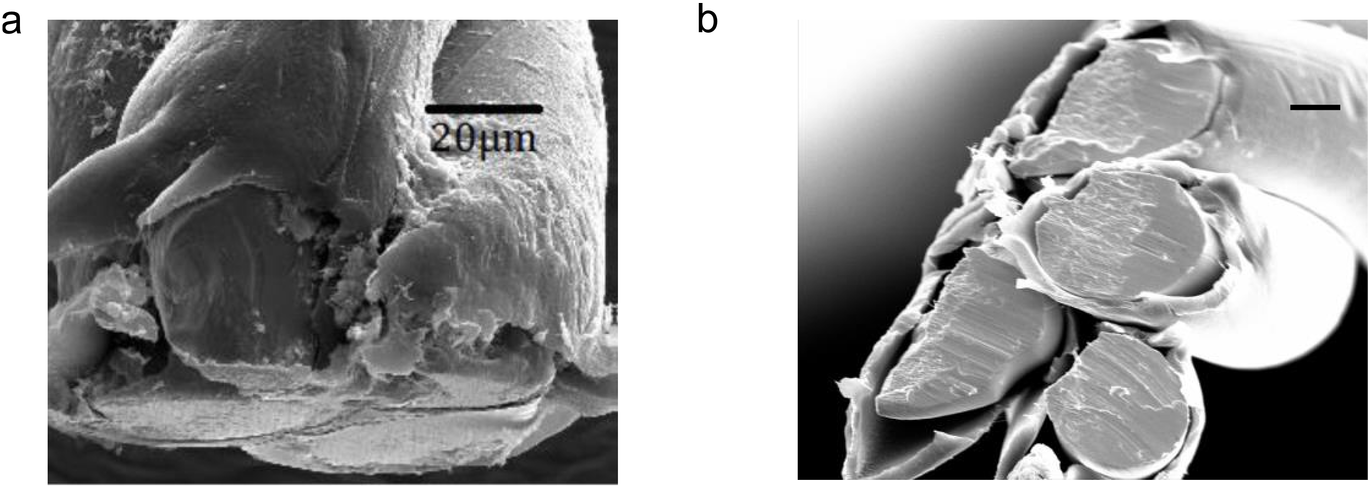
Polishing of tetrode tips. **a**, Tetrode tips can be polished with a circular grinder to 90 degrees after cutting. **b**, Empirically speaking, however, the polished tetrodes did not perform better than tetrodes simply cut with sharp carbide scissors at 45 degrees. We believe that the macroscopic surface roughness of the cut unpolished surface may be contributing to carbon nanotube adhesion. Scale bar: 10 μm.

**Supplementary Figure 8.**
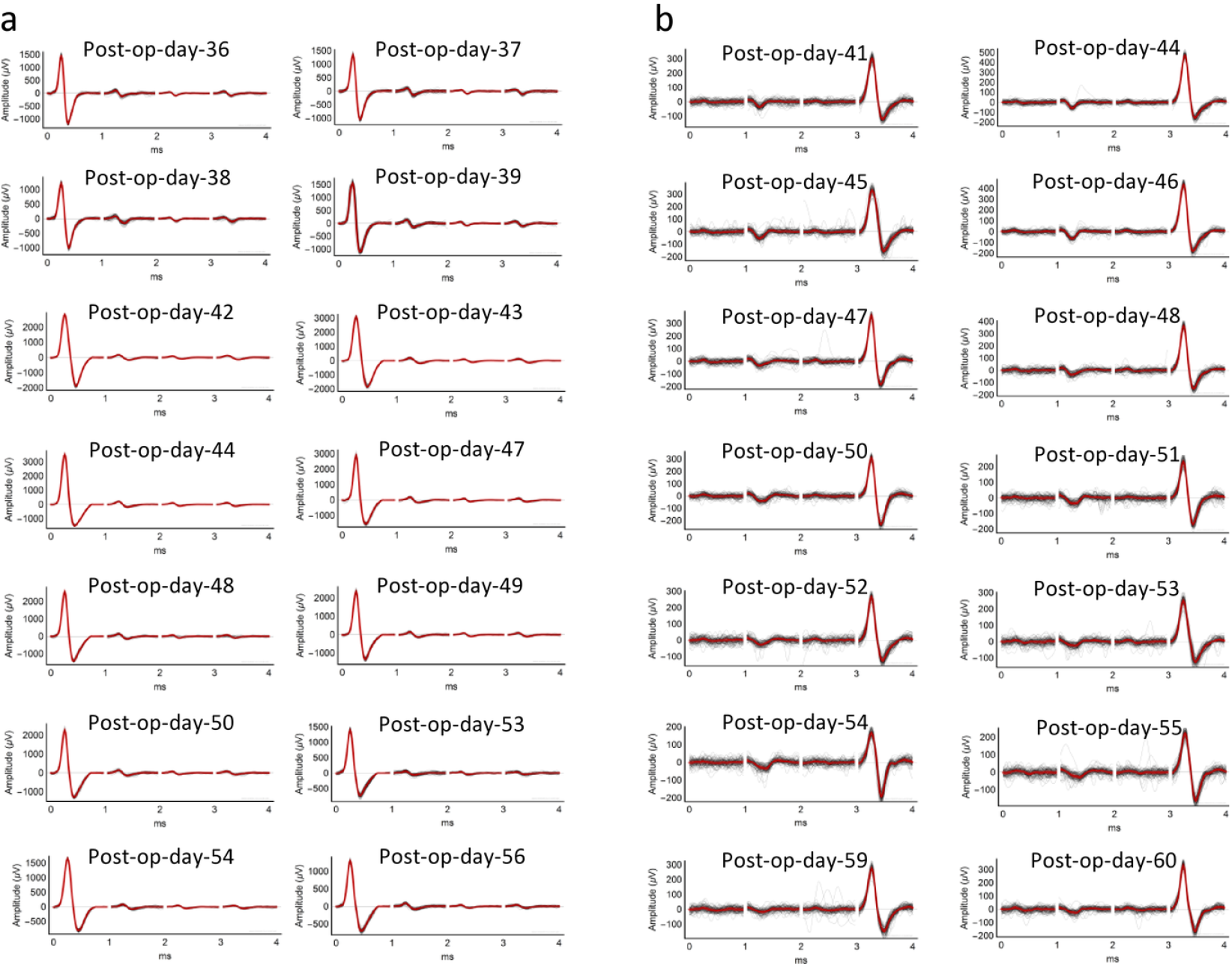
Single units can be recorded over weeks in multiple areas. Daily cluster waveforms of data shown in Fig. 5. **a** corresponds to panels a, c, e in Fig. 5. **b** corresponds to panels b, d, and e in Fig.5.

**Supplementary Figure 9.**
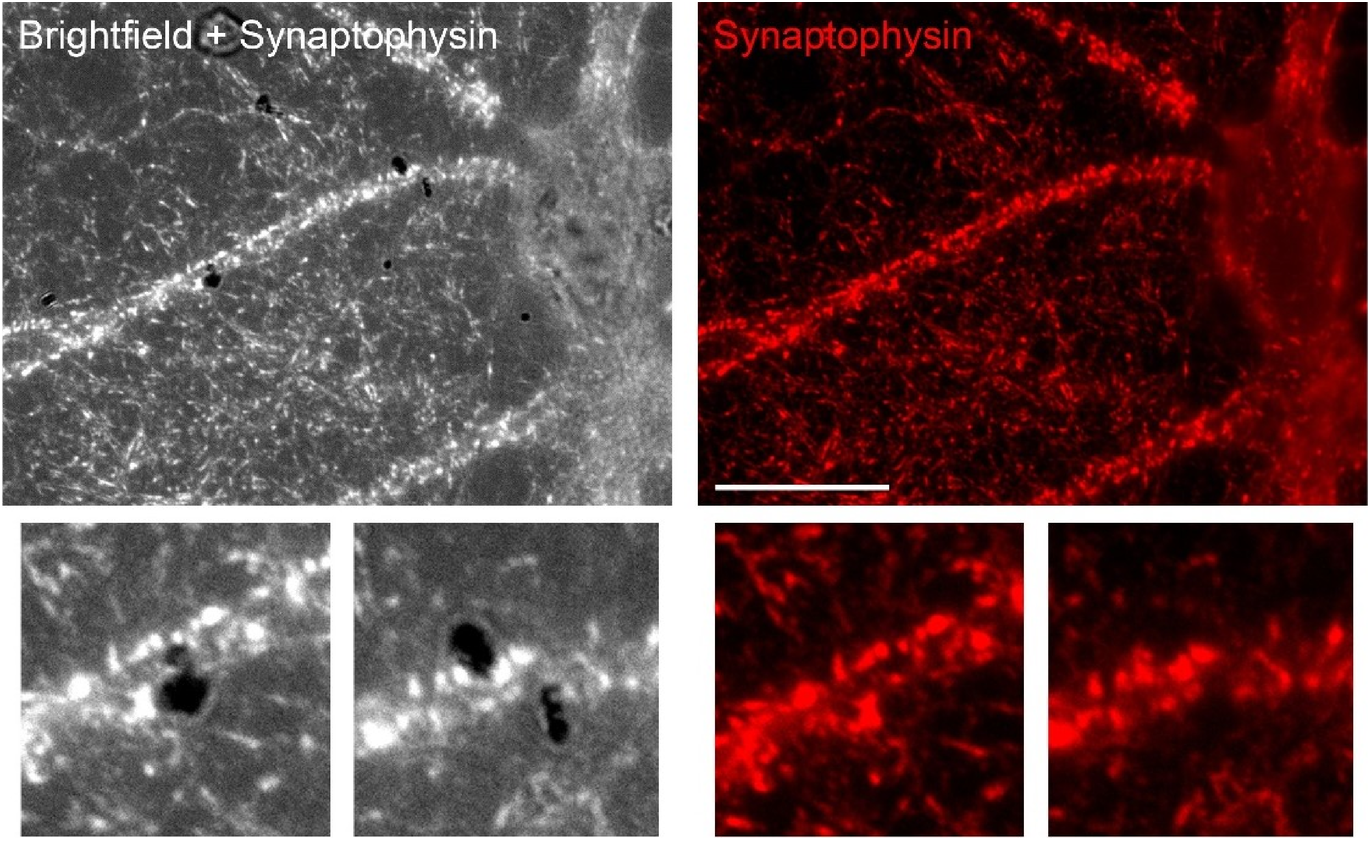
Cell morphology and synaptic contacts in neuronal cultures exposed to carbon nanotubes. After long-term incubation with carbon tubes (see Methods), hippocampal neurons were fixed and stained with an antibody against the presynaptic marker synaptophysin 1 (red) at 17 days in vitro. Intact cell morphology and the presence of dark precipitates of carbon nanotube were confirmed under combined brightfield-fluorescent illumination. Scale bar: 10 μm. Note that despite direct contact of several nanotubes, dendrites developed normally. Synapse number and fluorescent puncta size are grossly normal. Such nanointerfaces such as carbon nanotubes are known to support biocompatibility and biocontact ^43,44,45^.

## Notes

#### Summary of Updates

emphasizing novelty

